# Sustained organic loading disturbance favors nitrite accumulation in bioreactors with variable resistance, recovery and resilience of nitrification and nitrifiers

**DOI:** 10.1101/605733

**Authors:** E. Santillan, W. X. Phua, F. Constancias, S. Wuertz

## Abstract

Sustained disturbances are relevant for environmental biotechnology as they can lead to alternative stable states in a system that may not be reversible. Here, we tested the effect of a sustained organic loading alteration (food-to-biomass ratio, F:M, and carbon-to-nitrogen ratio, C:N) on activated sludge bioreactors, focusing on the stability of nitrification and nitrifiers. Two sets of replicate 5-liter sequencing batch reactors were operated at different, low and high, F:M (0.19-0.36 mgCOD/mgTSS/d) and C:N (3.5-6.3 mgCOD/mgTKN) conditions for a period of 74 days, following 53 days of sludge acclimation. Recovery and resilience were tested during the last 14 days by operating all reactors at low F:M and C:N (henceforth termed F:M-C:N). Stable nitrite accumulation (77%) was achieved through high F:M-C:N loading with a concurrent reduction in the abundance of *Nitrospira*. Subsequently, only two of the three reactors experiencing a switch back from high to low F:M-C:N recovered the nitrite oxidation function, with an increase in *Nitrobacter* as the predominant NOB, without a recovery of *Nitrospira*. The AOB community was more diverse, resistant and resilient than the NOB community. We showed that functional recovery and resilience can vary across replicate reactors, and that nitrification recovery need not coincide with a return to the initial nitrifying community structure.

## Introduction

Improving stability and optimizing performance of wastewater treatment processes are central tenets of environmental engineering and biotechnology, to help achieve the sustainable development goal of guaranteeing availability and sustainable management of water and sanitation for everyone^1^. In ecology, disturbances are believed to have direct effects on the stability of ecosystems by altering community structure and function^2^. Stability is multidimensional^3^, having several quantifiable aspects like resistance (ability to withstand disturbance), resilience (speed of recovery from disturbance), and recovery (ability to return to prior conditions after disturbance ceases)^4^. For engineered systems like activated sludge bioreactors, it is important to identify the effect of different disturbances on the microbial community structure so as to relate them to changes in process performance^5^. However, disturbance is also deemed to affect underlying mechanisms of community assembly^6^ which, if predominantly stochastic, could drive microbial communities to divergent trajectories in terms of composition and function^7^. Therefore, robust replication^8^ is valuable to assess the effect of disturbances in the stability of sludge bioreactors.

Sustained disturbances that impose a long-term continuous change of species densities through an alteration of the environment, also called press disturbances^9,10^, are relevant since they can lead a system to alternative stable states that may or may not be reversible in terms of both community composition and function^11^. Disturbances could be alterations in the environment that are not directly detrimental for organisms, but still provide opportunities for low abundance members within the community^12^. In bioreactors, a switch in the substrate feeding scheme employed could then elicit changes in community structure and function. Such disturbances can occur in the form of organic shocks within activated sludge^13–15^ and anaerobic reactor systems^16,17^. In wastewater treatment, the food-to-biomass ratio (F:M) or sludge loading rate is an important parameter as it determines the growth type and settleability of sludge microorganisms^18^. Yet, how press disturbances of F:M alterations affect the stability of microbial populations and specific functions in sludge bioreactors remains largely unknown.

Partial nitrification is an important stage in biological nitrogen removal from wastewater via nitritation/anammox^19^, for which the accumulation of nitrite is desired by promoting the growth of ammonia oxidizing bacteria (AOB) while suppressing nitrite oxidizing bacteria (NOB)^20^. Different strategies have been employed towards this goal, like varying temperature, dissolved oxygen (DO), pH, SRT, and substrate concentrations^21^. Variations in C:N are known to affect nitrification due to changes in competition for DO and space in biofilms^22^. However, substrate manipulation in particular has been reported to yield contrasting results in terms of C:N^23^. Low influent C:N values (2-3 mg 24 25-27 COD/mg TN) have been shown to prevent^24^, but also promote^25–27^ nitrite accumulation in different sludge systems. Other studies have reported nitrite accumulation at high (10 mg COD/mg NH_4_^+^-N)^28^ and also fluctuating influent C:N values (2.5-8 mg COD/mg NH_4_^+^-N)^29,30^. Additionally, although adjustments in C:N also impact F:M values by altering the available carbon for heterotrophic growth, covariations in F:M are rarely considered, and hence there is a knowledge gap about system performance and stability when both factors are controlled simultaneously. To uncover additional nitrite accumulation strategies, more research is needed to understand the effect of variations in influent C:N together with controlled variations of other important operational parameters like F:M.

The aim of this work was to test the effect of a press disturbance of doubling F:M and C:N (henceforth termed F:M-C:N) values in a set of replicated lab-scale sludge bioreactors after an acclimation period (Fig. 1). The focus was on the stability of the nitrite oxidation function and the nitrifying microbial community. Function dynamics were monitored throughout the study by periodic analysis of reactor effluent, as well as detailed temporal studies of reactor cycles at seven different time points. Nitrite accumulation was also tracked due to its relevance for practical applications. Changes in composition of nitrifying organisms and genes involved in nitrification were assessed by metagenomics and 16S rRNA gene amplicon sequencing. The resistance of nitrite oxidation and specific nitrifier abundance was quantified and evaluated by monitoring their transition to a different steady state after the disturbance. We further tested recovery and resilience by shifting the F:M-C:N back to the original pre-disturbance state.

**Fig. 1.**
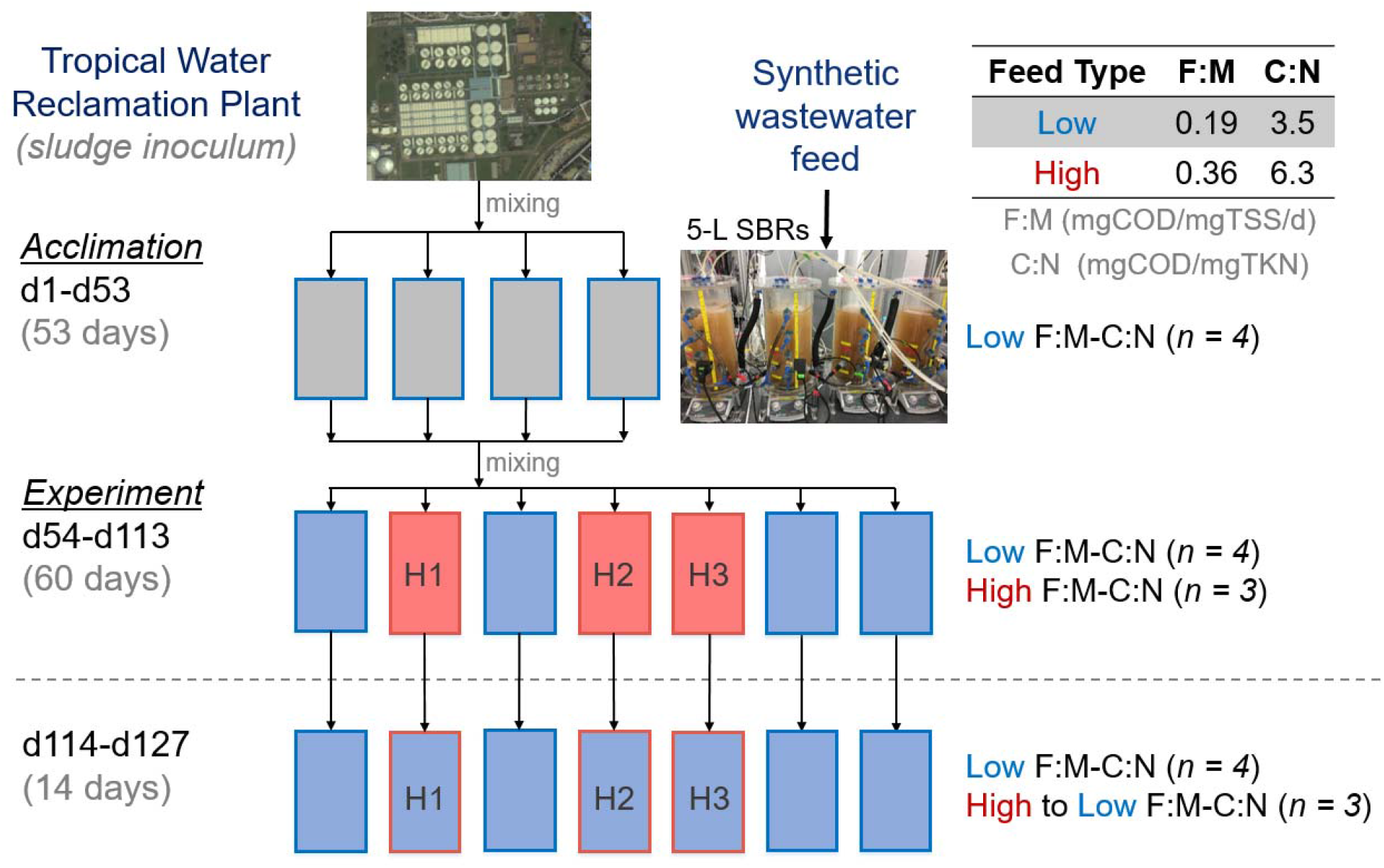
Schematic representation of the experimental design. F:M, food-to-biomass ratio; C:N, carbon-to-nitrogen ratio in the feed. Rectangles represent independent replicate 5-L SBRs. Phases: acclimation (grey), low F:M-C:N (blue, undisturbed), high F:M-C:N (red, press disturbed). Dashed line indicates removal of disturbance through the shift from high to low F:M-C:N.

## Results

### Dynamics in bioreactor performance

During the acclimation phase, the F:M-C:N values were maintained at 0.21 (mg COD/mg TSS/d) and 3.5 (mg COD/mg TKN), respectively (Table 1). Ammonium concentrations in the effluent decreased gradually while nitrate concentrations increased (Fig. 2). Sludge related parameters like settleability (SVI) and biomass fraction (VSS:TSS) varied during this acclimation period (Fig. S1). Most of these variations decreased after 30 d and trends were stable after 45 d. During the disturbance phase of the study (d54 onwards), sludge was wasted more often to better control the TSS and thus the F:M (Fig. S1), which is the reason why the SRT for the low F:M-C:N reactors is lower than during the acclimation phase. The average F:M and C:N values for the low F:M-C:N reactors were similar to those during the acclimation phase (0.19 and 3.5), while the ones for the high F:M-C:N reactors were controlled to be almost double (0.36 and 6.3). As expected (details in supplementary information), controlling for a higher F:M resulted in a lower solids residence time (SRT) in reactors subjected to this treatment (Table 1). This period showed a clear distinction between high and low F:M-C:N reactors in terms of nitrification (P < 0.014, Table S1), with the high F:M-C:N reactors displaying nitrite accumulation with high NO_2_^-^-N effluent concentrations (Fig. 2). To ensure that the partial nitrification was due to different F:M-C:N values and not a lack of available dissolved oxygen, we increased the aeration rate from 1 to 4 L min^-1^ from d97 onwards without observing significant changes in NO_2_^-^-N and NO_3_^-^-N effluent compounds. The last two weeks of the study involved shifting operational parameters in the high F:M-C:N reactors to match those of the low F:M-C:N ones (Table 1). During this period a transition towards recovery of the nitrite oxidation function was observed (P > 0.43), with high variability of effluent NO_2_^-^-N and NO_3_^-^-N across reactors (Fig. 2).

**Fig. 2.**
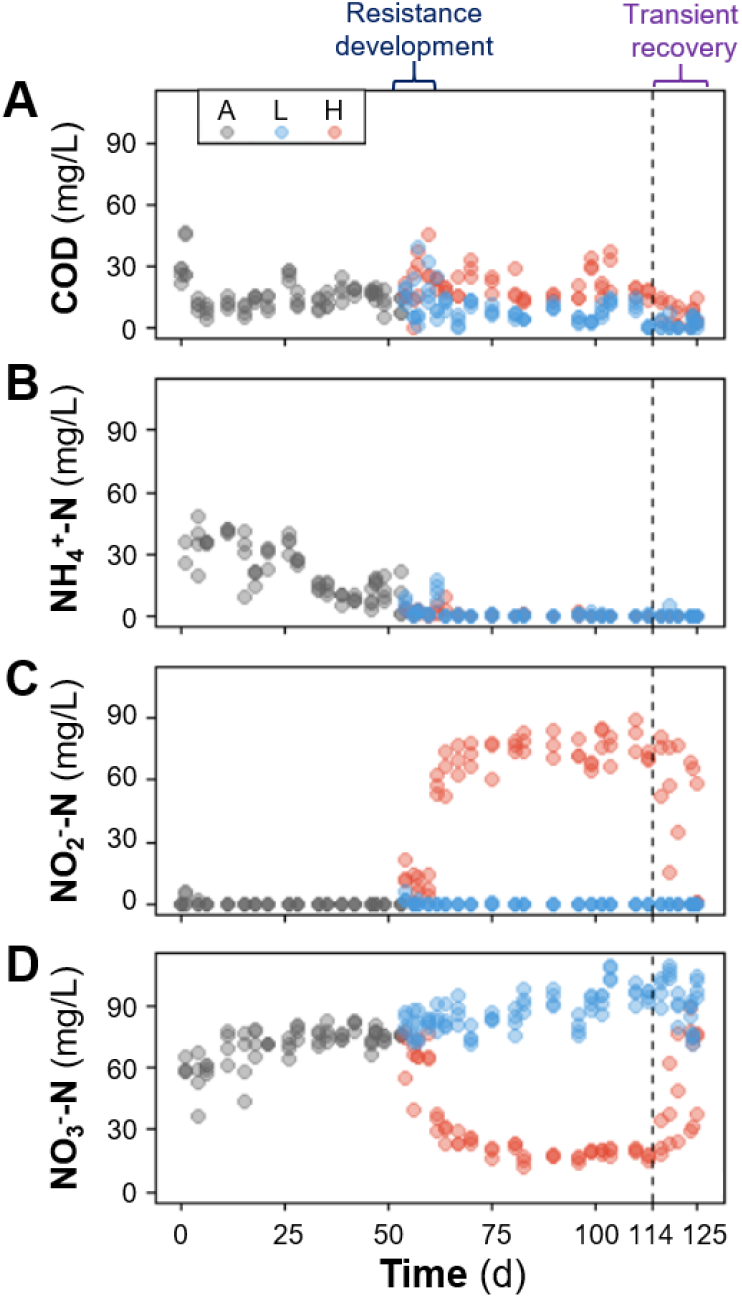
Temporal average effluent concentrations in mg/L of (**A**) soluble COD, (**B**) NH_4_^+^-N, (**C**) NO_2_^-^-N and (**D**) NO_3_^-^-N. Phases: A, acclimation (n = 4); L, low F:M-C:N (n = 4); H, high F:M-C:N (n = 3). Vertical dashed line indicates the shift from high to low F:M-C:N. Periods of functional resistance development and transient recovery are indicated by curly brackets.

**Table 1.** Influent synthetic wastewater characteristics and reactor operational parameters per phase.

| Day | Phase<sup>†</sup> | n | COD*<br>[mg/L] | TKN<br>[mg/L] | C:N<br>[mg COD/mg TKN] | TSS<br>[mg/L] | F:M<br>[mg COD/mg TSS/d] | SRT<sup>#</sup><br>[d] | VSS:TSS<br>[%] |
| --- | --- | --- | --- | --- | --- | --- | --- | --- | --- |
| **1-53** | Acclimation | 4 | 374 (106) | 105 (27) | 3.5 (0.7) | 1934 (502) | 0.21 (0.08) | 11.6 (0.4) | 92.0 (2.3) |
| **54-127** | Low F:M-C:N | 4 | 323 (24) | 92 (3.6) | 3.5 (0.3) | 1727 (251) | 0.19 (0.05) | 7.9 (0.2) | 96.1 (1.6) |
| **54-113** | High F:M-C:N<sup>‡</sup> | 3 | 629 (67) | 100 (19) | 6.3 (0.9) | 1943 (476) | 0.36 (0.11) | 5.1 (0.9) | 95.6 (1.4) |
| **114-127** | High to low F:M-C:N<sup>‡</sup> | 3 | 326 (19) | 90 (2.2) | 3.6 (0.3) | 1774 (256) | 0.19 (0.06) | 7.7 (0.6) | 95.4 (1.8) |

\*Average values, including standard deviation of the mean (s.d.m.) in parentheses.
<sup>†</sup> Each phase operated independent 5-L reactors. Samples were generated 2-3 times per week.
<sup>‡</sup> These two phases involved the same three reactors, where the F:M-C:N was changed from high to low on d114.
<sup>#</sup> Aerobic SRT, based on the aeration period (61.8%) of each cycle.

### Dynamics in nitrification and nitrifiers

The acclimation phase (d1-d53) displayed negligible nitrite accumulation at 0.3% (± 1.5%). Low F:M-C:N reactors had zero percent nitrite accumulation after d61 and only 1.1% (± 2.0%) during the first week (d54-d60). Conversely, high F:M-C:N reactors showed an initial transient nitrite accumulation of 18% (± 21%) on d54-d60, which subsequently increased and stabilized at 77% (± 6.0%) during the d61-d113 period. Finally, after shifting from high to low F:M-C:N conditions, nitrite accumulation decreased to 55% (± 29%) in the first week (d114-d120), and all the way to zero in the second week (d121-d127) for two of the three reactors (Fig. 2).

Among nitrifiers, the three most abundant bacterial genera detected through both metagenomics and 16S rRNA gene amplicon sequencing were *Nitrospira*, *Nitrosomonas* and *Nitrobacter*. During the disturbance phase, *Nitrospira* abundance was strongly reduced in high F:M-C:N reactors (to about 0.02%), while *Nitrosomonas* remained at around half the abundance levels observed for low F:M-C:N reactors (Fig. 3). This coincided with the accumulation of nitrite in high F:M-C:N reactors during the d54-d113 period of the study (Fig. 2). Similar patterns of nitrifier abundance across low and high F:M-C:N replicates were observed for both metagenomics and 16S rRNA gene amplicon sequencing datasets (Fig. 3). Further, relevant genes involved in nitrification (*amo*, *hao* and *nxr*) displayed a reduction in transcripts per million (tpm) across high F:M-C:N reactors compared to the low F:M-C:N reactors (Fig. 5). All the aforementioned differences in effluent values as well as nitrifier and nitrification gene abundances were statistically significant from d75 onwards (Table S1), until the disturbance was removed on d114.

**Fig. 3.**
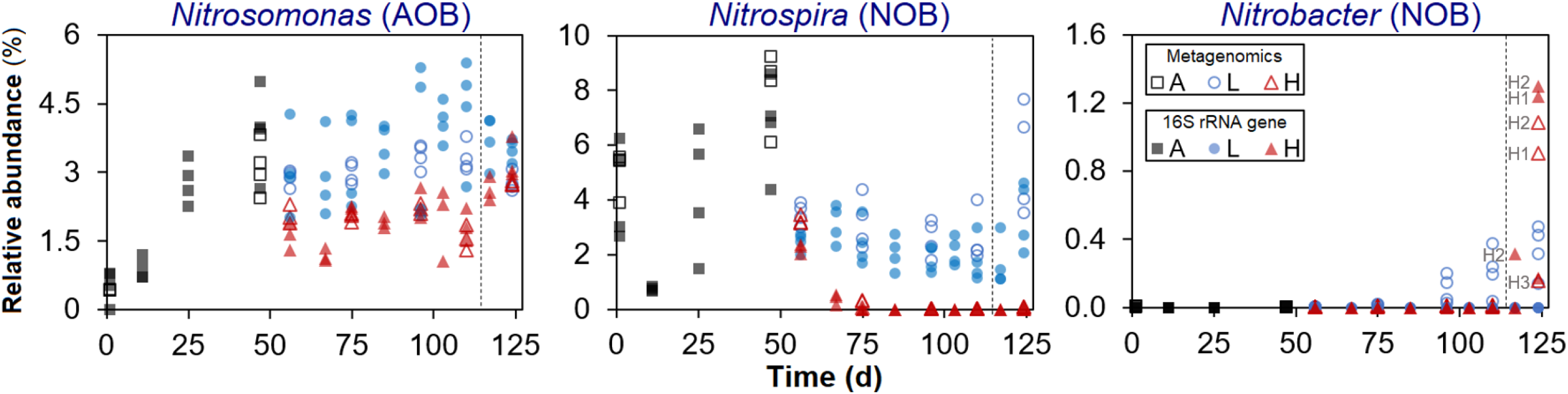
Temporal relative abundance of main nitrifier genera in each reactor. Phases: A, acclimation (n = 4); L, low F:M-C:N (n = 4); H, high F:M-C:N (n = 3). Vertical dashed line indicates the shift from high to low F:M-C:N. Closed symbols display 16S rRNA gene amplicon ASV data and open symbols shotgun metagenomics (summarized reads) data. Left, *Nitrosomonas*; centre, *Nitrospira*; right, *Nitrobacter*. There was variable NOB resilience as evident from the increase in *Nitrobacter* in two (H1 and H2) out of three reactors once the disturbance had been removed.

**Fig. 5.**
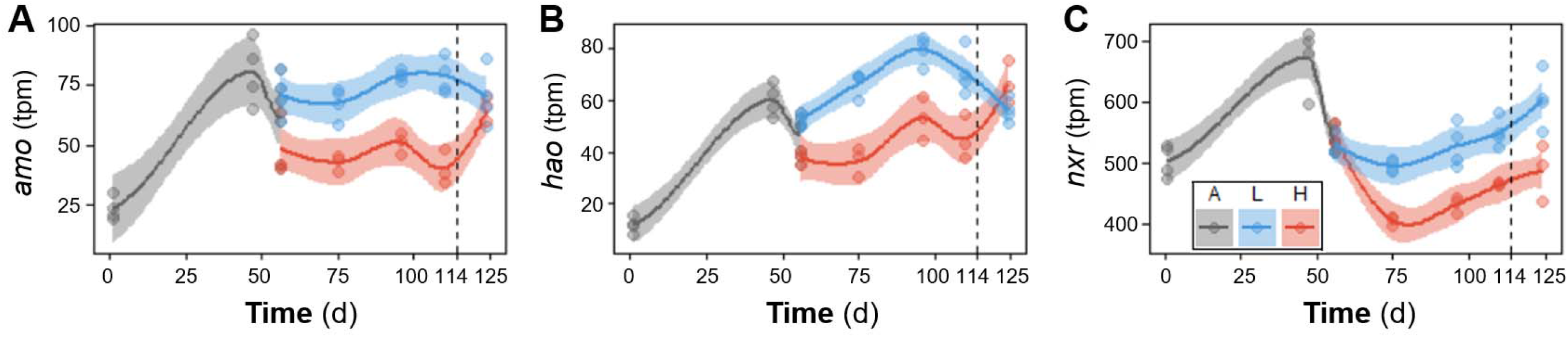
Temporal dynamics of nitrification genes *amo* (**A**), *hao* (**B**) and *nxr* (**C**), measured in transcripts per million (tpm) from assembled metagenomics data. Each point represents a different reactor on a given day. Phases: A, acclimation (n = 4); L, low F:M-C:N (n = 4); H, high F:M-C:N (n = 3). Vertical dashed line indicates the shift from high to low F:M-C:N. Lines refer to polynomial regression fitting, while shaded areas represent 95% confidence intervals.

Following the shift from high to low F:M-C:N it was *Nitrobacter* that rose to be the dominant NOB instead of *Nitrospira*, but only in two of three reactors (Fig. 3). Variations in performance among replicate reactors were also evident from cycle study profiles before (d110) and after (d124) the shift in feeding regime for the high F:M-C:N reactors (Fig. 4). Two weeks after the change, only two high to low F:M-C:N reactors displayed nitrite oxidation profiles similar to those of the low F:M-C:N reactors. The reactors that recovered functionality were the same as those that registered around 1% of *Nitrobacter* abundance (Fig. 3). By the end of the study (d124), abundances of *Nitrosomonas* as well as the tpm of *amo* and *hao* genes in the high to low reactors recovered to values that were not significantly different from the low F:M-C:N reactors (Figs. 3 and 5, Table S1).

**Fig. 4.**
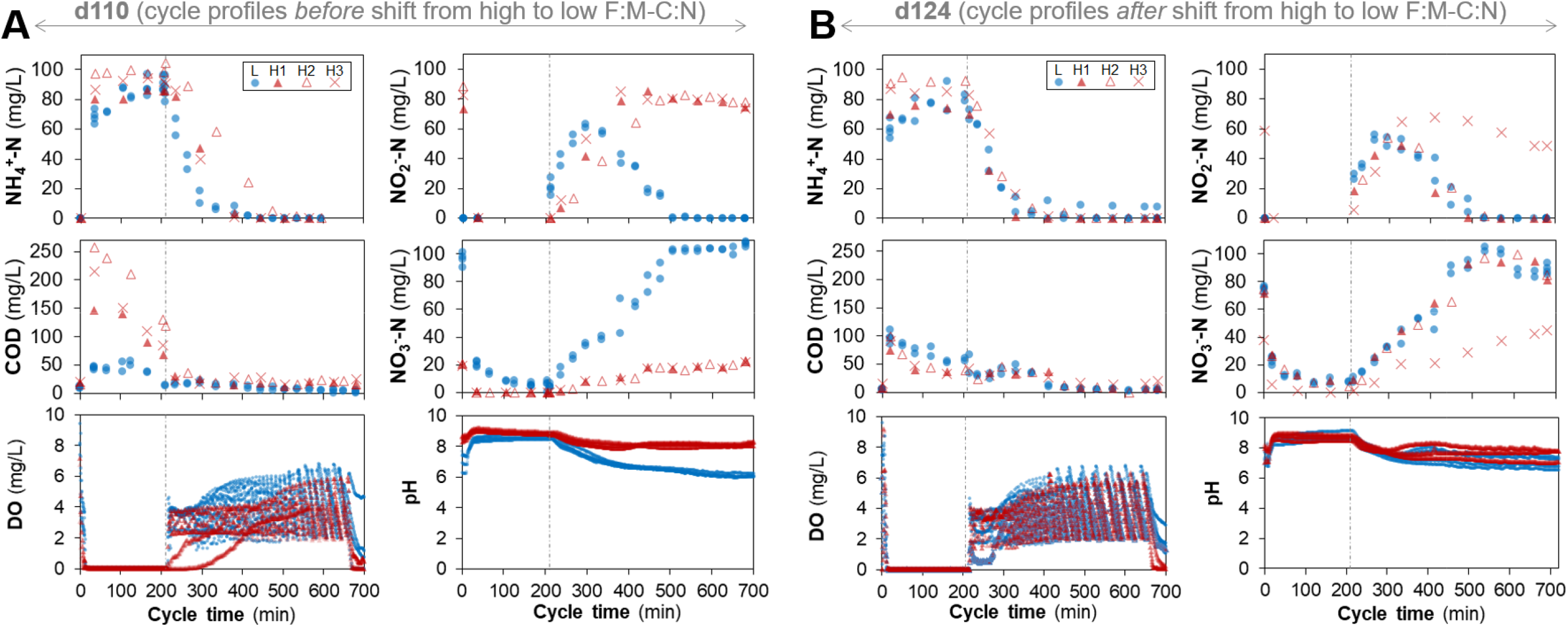
Chemical profiles during a full intermittently-aerated SBR cycle lasting 12 hours. Concentrations are shown for all reactors on (**A**) day 110 and (**B**) day 124, showing profiles before and after high to low F:M-C:N changes. L, low F:M-C:N reactors (n = 4); H, high F:M-C:N reactors (n = 3). Vertical dashed dotted line indicates the start of the aerobic stage in each cycle. Right panels (**B**) show that two (H1 and H2) out of three reactors fully recovered the nitrite oxidation function on d124.

Stability metrics for each of the high F:M-C:N replicate reactors (Fig. 6) showed a temporal reduction in resistance for both the nitrite oxidation function and the abundance of *Nitrospira*. Removal of the disturbance was followed by a temporal recovery of the nitrite oxidation function across all replicates. At the end of the study, this function had been completely restored for the two reactors (H1 and H2) that also displayed the highest nitrite oxidation resilience, as well as an overcompensation-recovery (c > 0) in *Nitrobacter* abundances. Overall, the ammonia oxidizer *Nitrosomonas* displayed higher resistance and resilience values than the nitrite oxidizer *Nitrospira*, which did not recover even after the disturbance had been removed (Fig. 6).

**Fig. 6.**
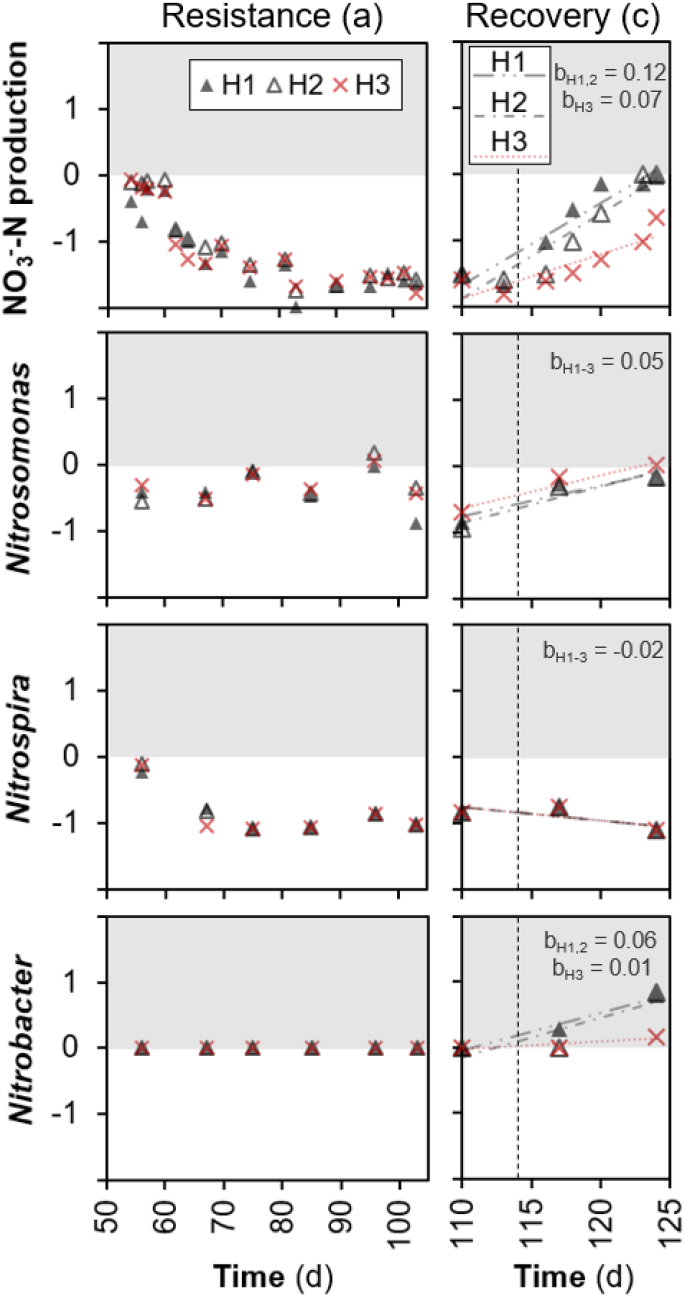
Stability metrics of resistance (a), resilience (b) and recovery (c) for the nitrite oxidation function and nitrifiers abundances across disturbed replicate reactors, adopted from^4^. Vertical dashed line indicates removal of disturbance. Dotted and dash-dotted lines represent linear regression fitting per replicate, with the slope (b) being the measure of resilience. Benchmarks: a = 0, maximum resistance; a < 0, low resistance through underperformance; b > 0, recovery; b <0, further deviation from undisturbed control; c = 0, maximum recovery; c < 0, incomplete recovery; c > 0, overcompensation.

The higher resolution of 16S rRNA gene amplicon sequencing allowed us to taxonomically identify four amplicon sequence variants (ASVs) as *Nitrospira*, 21 as *Nitrosomonas*, and one as *Nitrobacter* (Fig. S2). From these, only two ASVs were identified at the species level. The genus *Nitrospira* was dominated by the *N. defluvii* species, with the other three ASVs detected only at low abundances across the low F:M-C:N reactors after d97, the day the aeration rate was increased. The dominant *Nitrosomonas* ASVs during the disturbance phase were different from the initial ones. Furthermore, *N. europaea* and *Nitrosomonas* ASV-70 saw their relative abundances increasing with time across high F:M-C:N reactors (Fig. S2).

## Discussion

### Disturbance leads to stable nitrite accumulation unveiling system’s resistance

Sludge acclimation served to stabilize important functions like complete nitrification, organic carbon removal, and sludge settling capacity across reactors (Fig. 2, Fig. S1). Ecosystem functionality in terms of COD removal, ammonium removal, and complete nitrification was optimal and stable for the low F:M-C:N reactors, particularly towards the end of the study (Fig. 2). In contrast, press disturbed reactors (high F:M-C:N) experienced a marked reduction in nitrate production. The shift in function was not immediate, as the first seven days showed transient nitrite accumulation, which was variable among replicates, before stabilizing with little within-treatment variability for the remainder of the disturbance phase. Nitrite oxidation was not completely inhibited because 23% of the NO_X_^-^-N products still consisted of nitrate during the d61-d113 period. In addition to the calculated resistance metrics (Fig. 6), this percentage also shows the degree to which this particular function was insensitive to disturbance^32^. The higher variability in function among replicates during the first seven-day transition period could have been due to stochastic effects initially triggered in ecosystems after a disturbance^6^. Since the disturbance was sustained in this case, selective pressure likely promoted deterministic mechanisms resulting in a less variable function over time (Fig. 2). This is similar to what has been previously reported for studies on sludge bioreactors under sustained 3-chloroaniline disturbance^7^.

### Dynamics of nitrifiers during the disturbance phase

Understanding community and activity dynamics of nitrifying bacteria is essential for improving design and operation of wastewater treatment biological processes^33^. The changes in abundance of different *Nitrosomonas* ASVs suggested a succession of organisms within this genus (Fig. S2). Additionally, through the use of exact sequence variants^7^ we observed that *Nitrosomonas* was the most diverse nitrifying genus with 21 different ASVs being detected (Fig. S2). The reduction in *Nitrospira* abundances in the high F:M-C:N reactors (Fig. 3) could have been due to competition for DO with heterotrophs and *Nitrosomonas* that possess a higher affinity for oxygen^34,35^. It is known that AOB have a higher oxygen affinity than NOB^36^. Still, the increase in aeration rates from d97 onward did not prevent nitrite accumulation, indicating that a low DO in the system was not the main reason behind our observations. DO concentrations higher than 1 mg/L are enough to achieve optimal nitrification performance^37^, which was the case for the reactors in this study (Fig. 4). However, nitrifying communities grow in stratified biofilms where AOB are located closer to the water interface and NOB are in the interior zone^38,39^. The concentration of oxygen deep inside a biofilm or floc is lower than in the mixed liquor. Moreover, stratification in AOB biofilms due to an increase in C:N has been reported^34^, highlighting that increases in biofilm thickness due to heterotrophic growth further reduce oxygen diffusion inside, which is detrimental to the growth of NOB. In our study, the period during the aerobic phase of a cycle when almost all ammonium had been removed and COD concentrations were either low or remained constant (around 400-500 min, Fig. 4) implies that heterotrophs and AOB were not competing with NOB for oxygen anymore. Although this should have provided sufficient oxygen to NOB to be active during the remainder of the aerobic phase, an increase in nitrate production was not observed. A possible reason for this could be nitrite accumulation, which was reported to be toxic to NOB at high concentrations^40^. Hence, the observed NOB suppression at high F:M-C:N could have been due to a combination of competition for DO with heterotrophs and *Nitrosomonas*, reduced oxygen diffusion into the nitrifier biofilm due to heterotrophic growth, and nitrite accumulation due to AOB activity.

AOB growth rates are normally higher than those of NOB at 30 °C, which implies that the SRT can be reduced to achieve partial nitrification^41^. In our study, increasing F:M while keeping TSS constant implied an SRT reduction of 35% in the high F:M-C:N reactors compared to the low F:M-C:N reactors. It is conceivable that part of the observed reduction in nitrifiers was due to washout given their low growth rates. It was suggested based on mathematical modelling that a reduction in SRT has a stronger effect on NOB than on AOB^42^. However, the aerobic SRT of 5.1 d used to operate the high F:M reactors is common in activated sludge processes performing complete nitrification^18,43^, and around the operating SRT of 5-6 d at the full-scale plant that provided the sludge inoculum for this study. SRT values of 4-8 d have been suggested as optimum for nitrification in practice^43,44^, while complete nitrification has been reported for SRT values as low as 2 d^45^. Thus, our observed changes in nitrifiers and nitrification function were driven by controlled changes in F:M and C:N values and not by washout of NOB due to a low SRT.

### Importance of parameter covariations for nitrification studies

We showed that a combined high F:M-C:N approach led to stable and reproducible nitrite accumulation (77%) after seven days of transition. A comparison with earlier studies where either F:M or C:N was the parameter of interest reveals inconsistent outcomes. For example, similar to our results, conditions at high F:M resulted in higher nitrite accumulation compared to low F:M in studies on full-scale sludge systems that focused on the effect of varying F:M directly^46^ or indirectly through changes in SRT^47,48^. Likewise, nitrite accumulation was found at high influent C:N (10 mg COD/mg NH_4_^+^-N) in a pilot-scale study using a completely stirred tank reactor^28^. However, contrary to our results, low influent C:N values (1-3 mg COD/mg NH_4_^+^-N) in high-strength industrial wastewaters were reported to yield partial nitrification in a review of full-scale anammox processes^41^. Also, nitrite accumulation was reported at low C:N values in studies using a 35-L SBR (~2 mg COD/mg TN)^25^, a lab-scale 6.3-L SBR (3.33 mg COD/mg NH_4_^+^-N)^27^, and a pilot-scale continuous-flow A/O/A reactor (3.19 mg COD/mg NH_4_^+^-N)^26^. On the other hand, there was complete nitrification without nitrite accumulation at low influent C:N (3 ± 1 mg COD/mg TN) in pilot-scale membrane batch reactors^24^, similar to our results. Increasing C:N molar ratios from 2 to 5 was shown to significantly reduce nitrification rates in a laboratory denitrification-nitrification system^49^. Moreover, nitrite accumulation was even reported for fluctuating low and high influent C:N values (2.5-8 mg COD/mg NH_4_^+^-N) in two different pilot-scale studies using SBRs^29,30^. These multiple conflicting findings in the literature suggest that other factors in addition to C:N variations affect nitrification. As changes in C:N also affect F:M values by altering the available carbon for heterotrophic growth, we recommend to evaluate these interconnected parameters simultaneously.

Further, control of TSS is critical to operational control strategies in practice, based on either F:M or SRT^43,44^. In our study design, doubling the amount of COD in the influent almost doubled both F:M and C:N values, as operational TSS and influent TKN were controlled to remain close to constant. Doubling the influent COD also doubled the OLR, and higher COD concentrations also increased the biomass produced per unit time; thus more sludge had to be wasted to keep the TSS constant, reducing the SRT as a consequence (details in supplementary information). This point serves to illustrate that important operational parameters like SRT, C:N and OLR also co-vary with F:M^43^; thus it is important to account for such covariations during experimentation with sludge bioreactor systems.

### Recovery and resilience of nitrite oxidation and nitrifiers

Since disturbances often occur in biotechnological systems, it is desirable to understand the mechanisms of recovery after disturbance^50^, for which experimental replication is important to ensure reproducibility^8^ and capture fluctuations in process instability^51^. Hence, we tested whether returning the press-disturbed reactors to their previous low F:M-C:N level would lead to recovery and resilience. The switch back had significant effects on nitrifier communities, the nitrite oxidation function and nitrification genes. Relative abundances of *Nitrosomonas* genera recovered quickly; yet high F:M-C:N reactors exhibited variable functional resilience because after ten days of returning to pre-disturbance conditions, only two of three reactors completely recovered the nitrification function. This inconsistency among independent replicate reactors could be due to the stochastic growth after disturbance typically associated with r-strategists^52^ and ruderal organisms^53^. According to the r/K ecological framework, early niche colonization stages should favour r-strategists, whereas K-strategies should prevail at a later stage when many organisms attempt to colonize^54^. In our study, the variable nitrification recovery after removing the high F:M-C:N disturbance seemed to be due to stochastic colonization by r-strategist NOB (*Nitrobacter*), replacing the function performed by K-strategist NOB (*Nitrospira*) before the disturbance. Besides having a fast growth rate^55^ and prevailing at alternating conditions^56^, *Nitrobacter* has been shown to thrive at higher DO^57^ and nitrite^46^ concentrations, which explains why its abundance increased in the previously high F:M-C:N reactors after the shift to low F:M-C:N conditions. However, as part of a secondary succession process after disturbance, it is possible that *Nitrospira* would have recovered as a dominant NOB if more time had been allowed in the study. Further, these reactors constituted a closed system, which implies that the *Nitrobacter* colonizers came from low-abundance seed-bank populations. Disturbance was shown to open niches for bacterial colonization in open sludge systems^58^. Here, we showed that recruitment of organisms from the existing seed-bank is also possible after disturbance. As immigration of nitrifiers into sludge systems has been shown to alter the local community composition and function^59,60^, studies of the effect of disturbance in closed sludge systems can help disentangle the effect of immigrating populations.

In this study, functional resilience differed from nitrifying community resilience. Reactors with recovered function after returning to low F:M-C:N conditions (Fig. 4) still remained distinct from the control reactors in terms of NOB composition (Fig. 3). The fact that an altered community can perform the same functions as the original one supports the idea of functional redundancy^61^. This finding is similar to what was found in a full-scale sludge system after a press disturbance^48^, but contrary to what was reported for lake microbial systems after a pulse disturbance^62^. These contrasting reports highlight the complexity of assessing disturbance-diversity-function relationships^63^, as they depend not only on the system assessed but also on the disturbance frequency (i.e., pulse or press). Finally, the variability in the recovery of the nitrification function and fluctuations in nitrifier populations could only be captured thanks to the replication employed in the design of our study. As we move towards stable operation for microbial resource management^50^, future disturbance studies on sludge bioreactors should be robustly designed to ensure process reproducibility and highlight operational ranges where functional variability can be encountered. Overall, this study exemplifies how controlled disturbance studies on sludge communities using parameters within the range of plant operation can lead to insights that are both ecologically and practically meaningful.

## Materials and Methods

### Experimental design

The study was conducted using seven 5-liter bioreactors inoculated with activated sludge from a water reclamation plant in Singapore and operated as sequencing batch reactors (SBR) on continuous 12-h cycles with intermittent aeration (Fig. 1). Initially, four reactors were acclimated to lab conditions and fed with complex synthetic wastewater for 53 days. The complex synthetic feed was adapted from Hesselmann *et al*.^64^. At the start of the experiment (d54), the sludge of the acclimation reactors was thoroughly mixed and redistributed across seven reactors. From these, four were randomly selected and designated as high F:M and C:N reactors, receiving double the carbon substrate in terms of chemical oxygen demand (COD) amount in its feed as a press disturbance for 60 days. One of these reactors broke prematurely, reducing the count to three replicates. The remaining four reactors were operated as before at low F:M and C:N. During the last two weeks of the study (d114-d127), the feed for the high F:M and C:N reactors was adjusted to equal that of low F:M and C:N reactors (Table 1). For the sake of simplicity, we refer to both F:M and C:N as F:M-C:N. Details about sludge inoculum, acclimation phase and complex synthetic wastewater preparation are available as supplementary information.

### Operational parameters

The reactor temperature was maintained at 30°C and sludge was continuously mixed with a magnetic stirrer. In each cycle, SBR phases were: 5 min feed, 200 min anoxic/anaerobic react, 445 min aerobic react, 50 min sludge settle, and 20 min supernatant drain. The DO concentration was controlled at 2 – 6 mg/L during the aerobic phase. The pH ranged from 6 – 9, owing to alkalinity provided in the feed. Two cycles per day corresponded to a hydraulic retention time (HRT) of 24 h. Effluent and influent compositions were measured 2-3 times per week in accordance with Standard Methods^65^. The targets were soluble COD, total alkalinity, and nitrogen species (ammonium, nitrite and nitrate) in the liquid phase using colorimetric tests and ion chromatography. Nitrite accumulation percentage in the effluent was calculated as the ratio of nitrite concentration and the sum of nitrate and nitrite concentrations. Total organic carbon and total Kjeldahl nitrogen (TKN) were also measured in the influent. To control the F:M, sludge biomass was measured as total (TSS) and volatile suspended solids (VSS) twice a week, after which sludge wastage was done to target 1500 mg/L of TSS. Sludge volume index (SVI) was calculated from the liquid and sludge volumes measured in the reactors after 30 min settling and the TSS values obtained in the same cycle. Microbial community function was also investigated in the form of intensive sampling (every 30 to 60 min) over seven 12-h cycle studies conducted throughout the experiment. Detailed equations and explanations for F:M, C:N and SRT calculations, as well as analytical methods, are available as supplementary information.

### Bioreactor arrangement

Each of the SBRs employed in this study was equipped with: a magnetic stir plate to ensure mixed liquor homogeneity, a pair of EasySense pH and DO probes with their corresponding transmitters (Mettler Toledo), a dedicated air pump, a dedicated feed pump, a solenoid valve for supernatant discharge, and a surrounding water jacket connected to a re-circulating water heater. The different portions of the cycle were controlled by a computer software specifically designed for these reactors (VentureMerger, Singapore).

### 16S rRNA amplicon sequencing and reads processing

Bacterial 16S rRNA amplicon sequencing was done in two steps (for details see Supplementary Information). Primer set 341f/785r targeted the V3-V4 variable regions of the 16S rRNA gene^66^. The libraries were sequenced on an Illumina MiSeq platform (v.3) with 20% PhiX spike-in and at a read-length of 300 bp paired-end. Sequenced sample libraries were processed following the DADA2 bioinformatics pipeline^67^ using the version 1.3.3 of the *dada2* R-package. DADA2 allows inference of exact amplicon sequence variants (ASVs) providing several benefits over traditional OTU clustering methods^31^. Illumina sequencing adaptors and PCR primers were trimmed prior to quality filtering. Sequences were truncated after 280 and 255 nucleotides for forward and reverse reads, respectively, length at which average quality dropped below a Phred score of 20. After truncation, reads with expected error rates higher than 3 and 5 for forward and reverse reads were removed. After filtering, error rate learning, ASV inference and denoising, reads were merged with a minimum overlap of 20 bp. Chimeric sequences (0.18% on average) were identified and removed. For a total of 104 samples, 19679 reads were kept on average per sample after processing, representing 49.2% of the average input reads. Taxonomy was assigned using the SILVA database (v.132)^68^. Adequacy of sequencing depth after reads processing was corroborated with rarefaction curves at the ASV level (Fig. S3).

### Metagenomics sequencing and reads processing

Libraries were sequenced in one lane on an Illumina HiSeq2500 sequencer in rapid mode at a final concentration of 11pM and a read-length of 250 bp paired-end. In total, around 325 million paired-end reads were generated, with 3.4 ± 0.4 million paired-end reads on average per sample (total 48 samples). Illumina adaptors, short reads, low quality reads or reads containing any ambiguous bases were removed using *cutadapt*^69^. High quality reads (91.0 ± 1.4% of the raw reads) were randomly subsampled to an even depth of 4,678,535 for each sample prior to further analysis. Taxonomic assignment of metagenomics reads was done following the method described by Ilott *et al*.^70^. High quality reads were aligned against the NCBI non-redundant (NR) protein database (March 2016) using DIAMOND^71^ with default parameters. The lowest common ancestor approach implemented in MEGAN Community Edition v.6.5.5^72^ was used to assign taxonomy to the NCBI-NR aligned reads with the following parameters (maxMatches=25, minScore=50, minSupport=20, paired=true). On average, 36.8% of the high-quality reads were assigned to cellular organisms, of which 98.4% were assigned to the bacterial domain. Adequacy of sequencing depth was corroborated with rarefaction curves at the genus taxonomic level (Fig. S3). Identification and quantification of genes involved in the nitrogen cycle was performed using SqueezeMeta pipeline^73^. Read pairs were co-assembled using Megahit^74^. Open reading frames (ORFs) were then predicted from contigs using Prodigal^75^. Functional annotation was performed using DIAMOND^71^ against the KEGG database^76^. Read mapping against contigs was then performed using Bowtie2^77^ in order to quantify the abundance of genes among the different samples, and transformed in transcripts per million (tpm) values^73^.

### Stability measures

Stability metrics of resistance, resilience and recovery were calculated for each replicate reactor at the disturbed high F:M-C:N level (H1, H2 and H3) as described by Hillebrand *et al*.^4^. Metrics were applied to the nitrite oxidation function, measured as NO_3_^-^-N production, and to the relative abundances of the main nitrifier genera *Nitrosomonas*, *Nitrospira* and *Nitrobacter*, detected via 16S rRNA gene amplicon sequencing. Resistance (a) was measured as the initial log response ratio of a parameter (Fi) from each disturbed replicate versus the average value of the undisturbed reactors: a = ln(F_i,H1-3_/F_i,L_). Recovery (b) was measured in the same way but as the final log response ratio, after disturbance was removed. Resilience (c) was measured as the slope of regression of the aforementioned log response ratio over the recovery time, following disturbance removal.

### Statistical analyses

Welch’s ANOVA was employed for univariate testing. All reported p-values were corrected for multiple comparisons using a False Discovery Rate (FDR) of 5%^78^. Regression analyses were performed in R v3.5.1 using the *ggplot2* package.

### Data availability

DNA sequencing data are available at NCBI BioProjects PRJNA559245. See supplementary information for details about sludge inoculum and acclimation phase, complex synthetic wastewater preparation, chemical analysis, calculation of parameters (F:M, C:N, SRT), DNA extraction and purification, and 16S rRNA gene and metagenomics library preparation and sequencing.

## Acknowledgements

This research was supported by the Singapore National Research Foundation and Ministry of Education under the Research Centre of Excellence Program. We thank D.I. Drautz-Moses for her support with the 16S rRNA gene amplicon and metagenomics library preparation and sequencing pipelines employed, and T.J. Qiang and N.A.B.A. Latiff for their assistance with molecular work. S.S. Thi and A.F.B.M. Batcha are acknowledged for their support with analytical chemistry equipment. We thank J. Coppens for insightful discussions on nitrification. E.S. was partially supported by a Fulbright Fellowship.

## Author Contributions

ES and SW conceived the study. ES designed the experiment. SW obtained the funding for the study. ES and WXP performed the experiments and conducted laboratory and molecular analyses (except library preparation and sequencing). ES performed the 16S rRNA gene bioinformatics analyses. FC performed the metagenomics bioinformatics analyses. ES interpreted the data, generated the results, and elaborated the main arguments in the manuscript. ES and SW wrote the manuscript.

