## Supplementary Information for "Sustained organic loading disturbance favors nitrite accumulation in bioreactors with variable resistance, recovery and resilience of nitrification and nitrifiers"

*Sludge inoculum and acclimation phase*

Sludge inoculum was collected from one of the activated sludge tanks of a water reclamation plant in Singapore, with a Modified Ludzack-Ettinger (MLE) process configuration. Operation parameters were: Q ≈ 200,000 $m^3$/d, T ≈ 30 °C, pH ≈ 6.7, total suspended solids (TSS) ≈ 1500 mg/L, hydraulic residence time (HRT) = 8 h, and solids residence time (SRT) = 5-6 d. Typical influent concentrations were: total Kjeldahl nitrogen (TKN) ≈ 49 mg/L and total chemical oxygen demand (COD) ≈ 320 mg $L^{-1}$. The plant receives a mix of residential, commercial and industrial wastewater as its influent, operating continuously at C:N ≈ 6.5 mg COD/mg TKN and F:M ≈ 0.21 mg COD/mgTSS/d. It had a removal efficiency of around 80% for N and 90% for COD. Activated sludge was collected in 20-L containers and immediately transported to the lab. The suspension was manually mixed by shaking the closed container thoroughly before transferring it to four 5-L sequencing batch bioreactors (SBRs) (Fig. 1). Prior to starting the acclimation phase, the sludge from these reactors was mixed three times by removing half the content of the vessels and redistributing it among other vessels to homogenize the sludge across replicate reactors. This same mixing procedure had to be repeated on days 6, 14 and 27, to reduce the effect of operational problems encountered in some reactors on those days that resulted in partial loss of sludge (discharge valve failure on d6 in R7, accidental discharge during the settling phase on d14 in R8, and sludge loss after probe cleaning on d27 in R7). This also helped maintain similarity among replicate reactors before the disturbance phase. Final mixing of sludge was carried out on d54 just at the beginning of the low and high F:M-C:N disturbance phases, where the sludge from the four acclimation reactors was homogenized and distributed across eight reactors for the experimental phase. The original design for this study included four replicates for the high F:M-C:N level (n = 4). However, one of the reactors suffered an air diffuser blockage on d64 that impeded proper aeration, affecting the performance of this reactor compared to others. For that reason this reactor was not included in the study, reducing the total number of replicates for the high F:M-C:N level to three. Detailed information on process performance and microbial community dynamics in the failed reactor can be found in chapter 5 of Santillan[1]. No problems were encountered in any of the remaining seven reactors during the experimental phase from d54 onwards.

*Bioreactor feeding and complex synthetic wastewater preparation*

SBR phases were: 5 min feed, 200 min anoxic/anaerobic react, 445 min aerobic react, 50 min sludge settle, and 20 min supernatant drain. Following the end of the aerobic phase in any given cycle, bioreactors had an idle time (50 min) to allow sludge settling, after which supernatant effluent was discharged (about half the working volume of each reactor, or 2.5 L), followed by the replacement of the same volume with synthetic wastewater during the feeding phase at the beginning of the next 12-h cycle.

The composition of synthetic wastewater in the bioreactor feed was adapted from Hesselmann *et al.*[2]. It contained the following compounds, expressed in mg/L in mixed liquor in each reactor right after feeding: yeast extract (20.3), soy peptone (26.8), meat peptone (27), casein peptone (40.2), sodium acetate anhydrous (115.4), dextrose anhydrous (92.3), urea (85.2), ammonium bicarbonate (76.7), ammonium chloride (144), sodium dihydrogen phosphate monohydrate (32.6), sodium phosphate dibasic dihydrate (5.9), calcium chloride dihydrate (50.5), magnesium sulfate heptahydrate (112.5), and sodium bicarbonate (372.7) which was added to replace the alkalinity consumed during nitrification. The medium also contained 0.5 mL/L of a trace element stock which contained (g/L) citric acid monohydrate (2.5), EDTA acid disodium salt dihydrate (0.6), hippuric acid (2), sodium molybdate dihydrate (0.12), potassium iodide (0.12), sodium tungstate dihydrate (0.12), boric acid (1), cobalt(II) chloride hexahydrate (0.12), copper(II) sulfate pentahydrate (0.24), manganese(II) chloride tetrahydrate (0.48), nickel(II) chloride hexahydrate (0.12), nitrilotriacetic acid trisodium salt monohydrate (1.44), iron(III) chloride hexahydrate (6), zinc sulfate heptahydrate (0.6).

The above values resulted in an average feed concentrations of 323 mg COD/L and 92 mg TKN/L in the mixed liquor (after feeding) for low F:M-C:N reactors. The high F:M-C:N reactors received double the amount of yeast extract (40.6), soy peptone (53.6), meat peptone (54), casein peptone (80.4), sodium acetate anhydrous (230.8), and dextrose anhydrous (184.6), which resulted in average feed concentrations of 629 mg COD/L and 100 mg TKN/L in the mixed liquor. The slight increase in TKN for high F:M-C:N reactors was due to the presence of organic N in the peptones and yeast extract, as we aimed to keep the mix of carbon sources equal for both low and high levels. Phosphate addition targeted a concentration in mixed liquor of 16 mg P/L to obtain a N:P of around 6.

*Chemical analysis*

Water quality parameters were measured in accordance with Standard Methods[3] and targeted COD (Standard Methods 5220 D) and nitrogen species (ammonium, nitrite, and nitrate ions) using spectrophotometric tests (Hach Company, Loveland, Colorado, USA) and Ion Chromatography (Standard Methods 4500-NH3 for ammonium; 4110 B for nitrate and nitrite). The COD measured was adjusted by subtracting the contribution of nitrite on the basis of 1.1 g COD/g $NO_2^-$-N to correct for nitrite interference. Total organic carbon (TOC) and total Kjeldahl nitrogen (TKN) were also measured in influent samples using a TOC-L analyzer (Shimadzu Corp., Kyoto, Japan). TSS, VSS and SVI measurements were done in accordance with Standard Methods[3]. Effluent samples were filtered through a 0.2-μm pore size filter and the filtrate was stored at 4°C for less than one week prior to chemical analyses.

*Operational parameter calculations*

Equations (1-5) below for operational parameters of food-to-biomass ratio (F:M), carbon-to-nitrogen ratio (C:N), hydraulic retention time (HRT), solids retention time (SRT) and organic loading rate (OLR), were based on Tchobanoglous *et al.*[4]:

$$F{:}M = \frac{Q \cdot S_0}{V \cdot X} \quad (1) \qquad C{:}N = \frac{S_0}{SN_0} \quad (2) \qquad HRT = \frac{V}{Q} \quad (3)$$

$$SRT = \frac{V \cdot X}{(Q - Q_W) \cdot X_e + Q_W \cdot X_r} \quad (4) \qquad OLR = \frac{Q \cdot S_0}{V} \quad (5)$$

where:

Q = flowrate, L/d

$Q_W$ = waste sludge flowrate, L/d

V = working volume, L

$S_0$ = influent organic carbon concentration (mg/L)

$SN_0$ = influent nitrogen as ammonium concentration (mg/L)

X = biomass concentration (mg/L)

$X_e$ = concentration of biomass in the effluent (mg/L)

$X_r$ = concentration of biomass in the return from clarifier (mg/L)

In this study enough settling time (50 min) was allocated at the end of each cycle before effluent discharge, thus the biomass content in the supernatant was negligible ($X_e \approx 0$). In practice, biomass is often measured as mixed liquor suspended solids, thus X = TSS. After two cycles (1 d) the total working volume of each reactor was replaced, therefore V/Q = 1 d. The influent organic carbon concentration was the COD in the mixed liquor after feeding ($S_0$ = COD). The influent nitrogen as ammonium concentration was the TKN in the mixed liquor after feeding ($SN_0$ = TKN). Sludge wastage was done twice a week for each reactor, in the same cycle where TSS was measured, and its biomass concentration was equal to that of the reactor ($X = X_r$). The volume of sludge wasted depended on the TSS value measured and was calculated to be such that, after feeding in the beginning of the next cycle, the biomass in the reactor would be TSS = 1500 mg/L. The waste sludge flowrate, $Q_W$ (L/d), was estimated for each reactor as the total sludge volume wasted in a reactor over a phase, divided by the total number of days of this phase. SRT calculations were adjusted to represent the aerobic fraction of the cycle (445 min during each cycle lasting 720 min) as required for SBRs[4].

Given the aforementioned considerations, equations (1-5) can be rewritten as follows:

$$F\!:\!M = \frac{COD}{TSS \cdot d} \quad (1') \qquad C\!:\!N = \frac{COD}{TKN} \quad (2') \qquad HRT = d \quad (3')$$

$$SRT = \frac{V}{Q_W} \cdot \frac{445\ min}{720\ min} \quad (4') \qquad OLR = \frac{COD}{d} \quad (5')$$

*Operational parameters covariations (F:M, SRT, C:N)*

The covariations among operational parameters that were described in the main text can be understood from equations (1'-5'). This study manipulated F:M by increasing influent COD while aiming to keep a constant TSS. More COD supports more biomass growth, which means that more sludge has to be wasted (higher $Q_W$) to keep TSS constant, thus the SRT decreases. Note that if TSS is not controlled, then F:M is not controlled either. This is why at constant TSS a higher F:M will always result in a lower SRT, and vice versa. Additionally, an increase in influent COD also increases C:N values, unless additional TKN is provided to proportionally compensate the COD change. In our study, such an adjustment would have implied very high influent nitrogen concentrations of 180 mg TKN/L, which would have confounded our observations. Therefore, we decided to allow the increase in F:M to occur concurrently with an increase in C:N in a controlled manner.

*DNA extraction and purification*

Aliquots of sludge samples were flash frozen in liquid nitrogen immediately after collection and stored at -80°C for a maximum of 12 months before molecular analysis. Genomic DNA was extracted from about 500 µL of sludge using the FastDNA Spin Kit for Soil and the FastPrep instrument (MP Biomedicals, Santa Ana, California, USA) with modifications to the manufacturer's protocol to increase DNA yield. The first modification involved performing four lysis cycles in the FastPrep Instrument instead of one, with two minutes of rest in between each cycle, during which the samples were placed on ice[5]. The second modification involved eluting DNA from the spin column using nuclease-free water (Qiagen, Venlo, Netherlands) that had been pre-heated to 55°C, followed by incubation of the columns in elution water at 55°C for five minutes before the final centrifugation. Extracted DNA was quantified using both NanoDrop 2000c and Qubit 3.0 fluorometer (both ThermoFisher Scientific, Waltham, Massachusetts, USA), and purified using the Genomic DNA Clean & Concentrator kit (Zymo Research Corp, Irvine, USA) following the protocol from the manufacturer.

*16S rRNA amplicon library preparation and sequencing*

For the first PCR stage, each reaction (25 µL) contained 12.5 µL of HiFi Hotstart Readymix (Kapa Biosystems), 9.5 µL of nuclease free water, 0.5 µL (each) of forward and reverse primers (10 µM) and 2 µL of DNA template (6 ng $\mu L^{-1}$). Primer set 341f/785r targeted the V3-4 variable regions of the 16S rRNA gene[6]. Thermocycler settings were: Initial denaturation at 95°C for 2 min, 30 cycles of 95°C for 30 s, 58°C for 15 s, 72°C for 30 s, and final elongation at 72°C for 2 min. PCR reactions were all run in duplicate and pooled subsequently. Amplicon libraries were purified using the Agencourt AMpure XP bead protocol (Beckmann Coulter). Library concentration was measured with Qubit 3.0 fluorometer (Thermo Fisher Scientific) and quality validated with a Tapestation 2200 (Agilent).

The second stage PCR (Indexing PCR) was performed according to the recommendations in Illumina's '16S Metagenomic Sequencing Library Preparation' application note. This step uses a limited 8-cycle PCR to complete the Illumina sequencing adapters and add dual-index barcodes to the amplicon target. Five microliters of the intermediate PCR product from the first stage were used as template for the indexing PCR and samples were amplified with 8 PCR cycles. Nextera XT v2 indices were used for dual-index barcoding to allow pooling of the amplicon targets for sequencing.

Finished amplicon libraries were quantitated using QuantiFluor dsDNA assay (Promega) and the average library size was determined on a Tapestation 4200 (Agilent). Library concentrations were then normalized to 4nM and validated by qPCR on a QuantStudio-3 system (Applied Biosystems), using the Kapa library quantification kit for Illumina platforms (Kapa Biosystems). The libraries were then pooled at equimolar concentrations and sequenced on an Illumina MiSeq platform (v.3) with 20% PhiX spike-in and at a read-length of 300bp paired-end. Sequencing was done at SCELSE's core sequencing facility. After bioinformatics processing, all 104 samples were rarefied to the one with the minimum number of reads passing the *dada2* pipeline (Fig. S3). Sample 29, corresponding to one of the high F:M-C:N reactors at d96, had to be re-sequenced due to initial low amplicon quality.

*Metagenomics library preparation and sequencing*

Prior to library preparation, the quality of the DNA samples was assessed on a Bioanalyzer 2100, using a DNA 12000 Chip (Agilent). Sample quantitation was carried out using Invitrogen's Picogreen assay. Library preparation was performed according to Illumina's TruSeq Nano DNA Sample Preparation protocol. DNA samples were sheared on a Covaris E220 to ~450bp, following the manufacturer's recommendation, and uniquely tagged with one of Illumina's barcodes to allow pooling of libraries for sequencing. The finished libraries were quantitated using Invitrogen's Picogreen assay and the average library size was determined on a Bioanalyzer 2100, using a DNA 7500 chip (Agilent). Library concentrations were then normalized to 4nM and validated by qPCR on a ViiA-7 real-time thermocycler (Applied Biosystems), using the KAPA Illumina Library Quantification Kit (Kapa Biosystems, Roche). The libraries were then pooled at equimolar concentrations and sequenced in one lane on an Illumina HiSeq2500 sequencer in rapid mode at a final concentration of 11pM and a read-length of 250 bp paired-end. Sequencing was done at SCELSE's core sequencing facility. After bioinformatics processing, all 48 samples were rarefied to have an even number of 1,661,886 total summarized bacterial reads per sample (Fig. S3). Genus-level relative abundance calculations were done by taking into account the summarized genus level reads and the total summarized bacterial reads per sample.

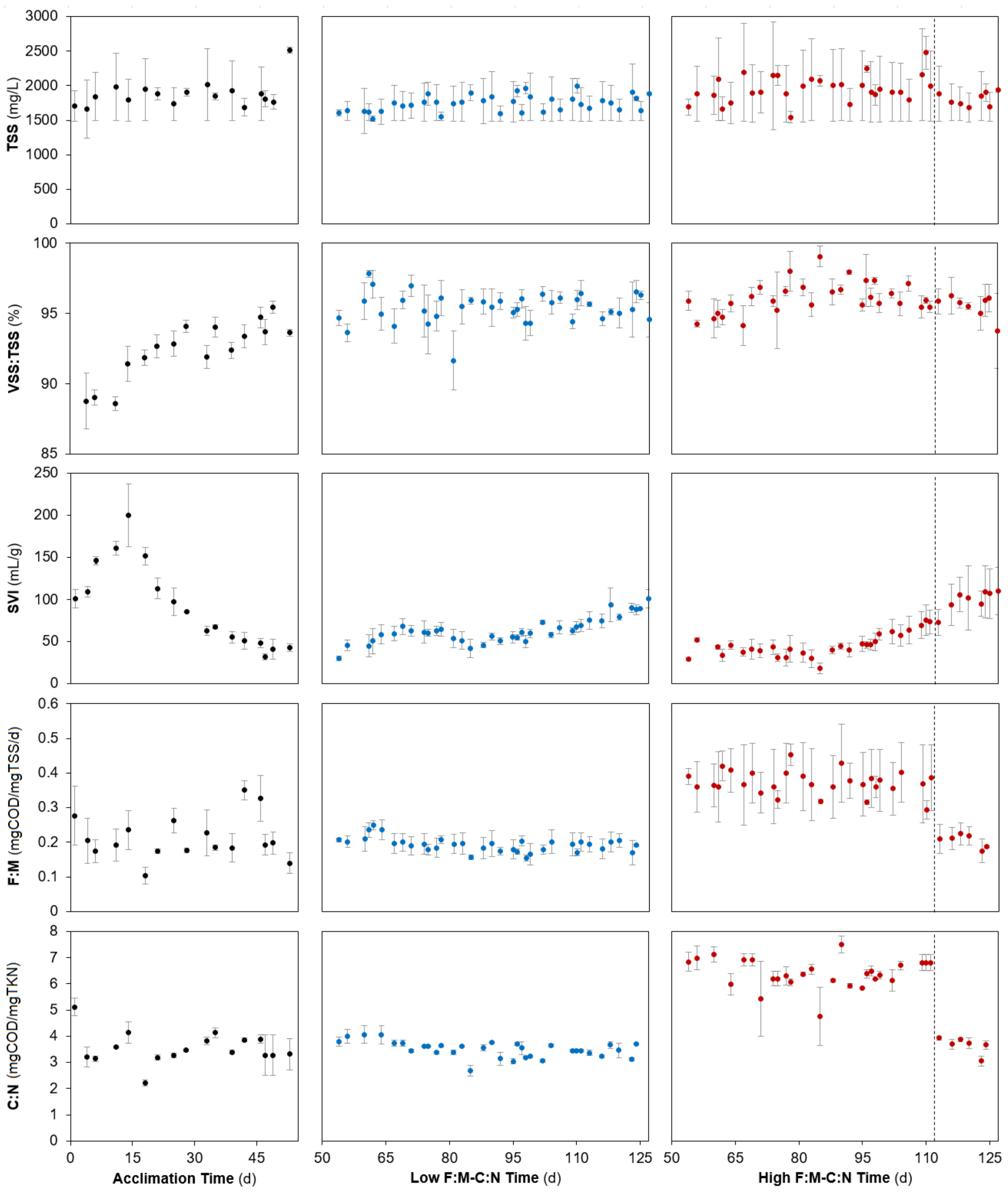


**Fig. S1.** Temporal average mixed liquor characteristics. Left, acclimation phase (black dots); centre, low F:M-C:N phase (blue dots); right, high F:M-C:N phase (red dots). Error bars represent one s.d.m. Vertical dashed line indicates the shift from high to low F:M-C:N. TSS, total suspended solids; VSS, volatile suspended solids; SVI, sludge volume index; F:M, food-to-biomass ratio; C:N, carbon-to-nitrogen ratio in the feed.

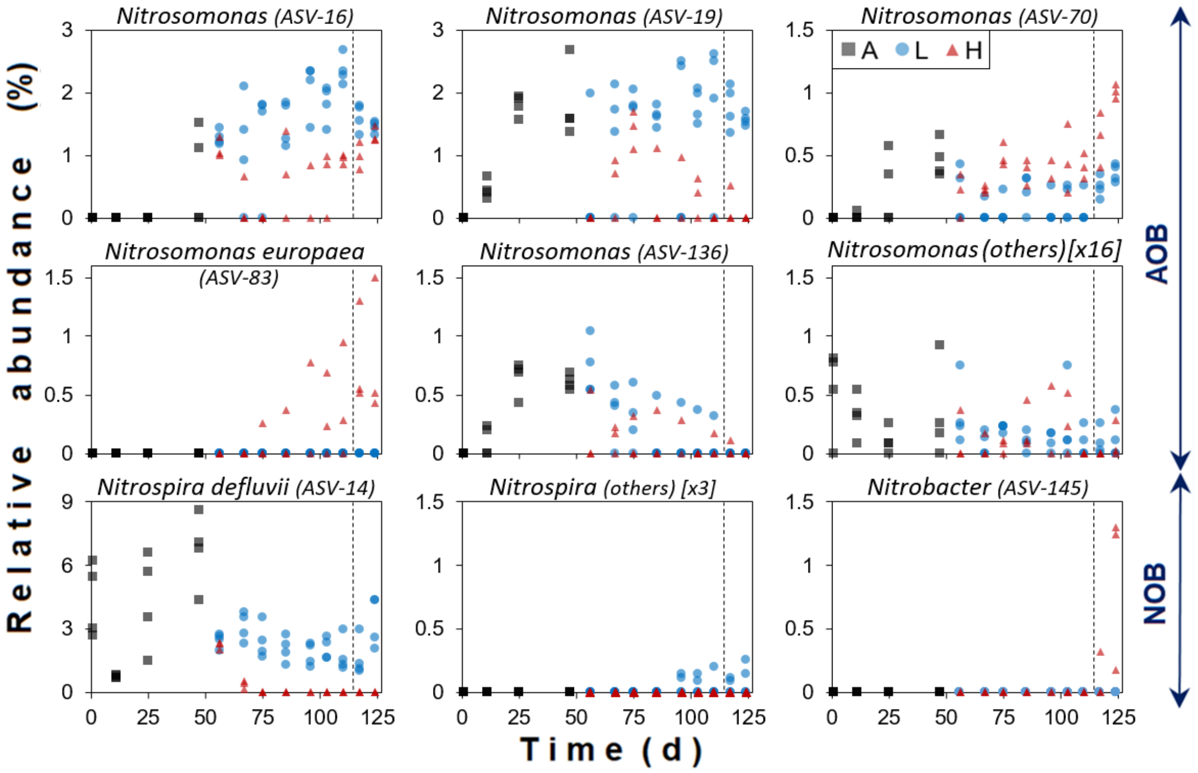


**Fig. S2.** Temporal relative abundance of ASVs assigned to nitrifier genera in each reactor. Phases: A, acclimation (n = 4); L, low F:M-C:N (n = 4); H, high F:M-C:N (n = 3). Vertical dashed line indicates the shift from high to low F:M-C:N. The abundance rank number (among all 1646 ASVs detected) is shown in parentheses. Brackets indicate the number of other additional lower abundance ASVs summed as 'others' within a panel. The higher number of ASVs detected suggests that AOB populations were more diverse than NOB populations.

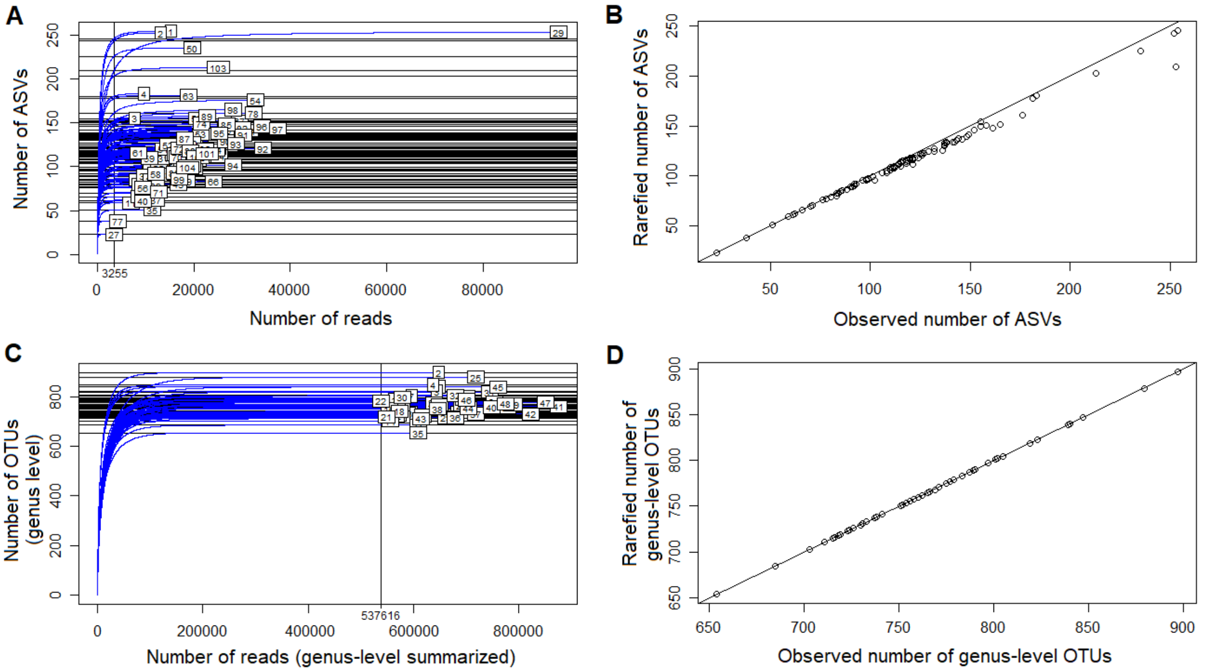


**Fig. S3.** Rarefaction plots for (**A-B**) 16S rRNA gene and (**C-D**) metagenomics datasets. Left panels, rarefaction curves in blue for (**A**) ASVs and (**B**) all genus-level OTUs. Numbers in boxes represent sample numbers. Vertical line indicates rarefaction level which corresponded to the sample with the minimum number of reads after bioinformatics processing. Right panels, rarefied versus observed number of (**B**) ASVs and (**D**) genus-level OTUs.

**Table S1.** P-values for Welch's t-tests between low (n = 4) and high (n = 3) F:M-C:N treatments over time for function, nitrifiers and nitrification genes abundances over time. P-values adjusted at a FDR of 5%.

| **d** | **Function[*]** | | | **Nitrifiers[†]** | | **Nitrification genes[‡]** | | |
|---|---|---|---|---|---|---|---|---|
| | **COD** | **$NO_2^-$-N** | **$NO_3^-$-N** | ***Nitrosomonas*** | ***Nitrospira*** | ***nxr*** | ***amo*** | ***hao*** |
| 56 | 0.900 | 0.450 | 0.177 | 0.014 | 0.419 | 0.419 | 0.110 | 0.007 |
| 75 | 0.024 | 0.014 | 0.000 | 0.007 | 0.018 | 0.000 | 0.006 | 0.006 |
| 96 | 0.051 | 0.006 | 0.000 | 0.006 | 0.013 | 0.018 | 0.007 | 0.028 |
| 110 | 0.039 | 0.008 | 0.001 | 0.005 | 0.024 | 0.014 | 0.007 | 0.029 |
| 124 | 0.075 | 0.457 | 0.434 | 0.900 | 0.024 | 0.044 | 0.548 | 0.177 |

[*] Tested effluent parameters in mg/L at the end of a bioreactor cycle (see values in Fig. 2).

[†] Tested relative abundances from metagenomics summarized reads data (see values in Fig. 3).

[‡] Tested transcripts per million from metagenomics assembled data (see values in Fig. 6).
